# A simple method to quantify protein abundances from one thousand cells

**DOI:** 10.1101/753582

**Authors:** Burcu Vitrinel, Dylan E. Iannitelli, Esteban O. Mazzoni, Lionel Christiaen, Christine Vogel

**Affiliations:** Center for Genomics and Systems Biology, Department of Biology, New York University, New York, NY, USA; Center for Developmental Genetics, Department of Biology, New York University, New York, NY, USA; NYU Neuroscience Institute,NYU Langone Medical Center, New York, NY, USA

## Abstract

The rise of single-cell transcriptomics has created an urgent need for similar approaches that use a minimal number of cells to quantify expression levels of proteins. We integrated and optimized multiple recent developments to establish a proteomics workflow to quantify proteins from as few as 1,000 mammalian stem cells. The method uses chemical peptide labeling, does not require specific equipment other than cell lysis tools, and quantifies >2,500 proteins with high reproducibility. We validated the method by comparing mouse embryonic stem cells and *in vitro* differentiated motor neurons. We identify differentially expressed proteins with small fold-changes, and a dynamic range in abundance similar to that of standard methods. Protein abundance measurements obtained with our protocol compare well to corresponding transcript abundance and to measurements using standard inputs. The protocol is also applicable to other systems, such as FACS-purified cells from the tunicate *Ciona*. Therefore, we offer a straightforward and accurate method to acquire proteomics data from minimal input samples.

## Introduction

Mass spectrometry is a highly sensitive technique capable of detecting peptides in the femtogram scale ^1^. However, to reach maximal protein identification and reproducibility, the quality and quantity of the sample material holds a great importance. Therefore, standard proteomics analysis uses large numbers of cells, estimated at 10^6^ to 10^8^ ^2^ to identify a few thousand proteins in a single-shot tandem mass spectrometry experiment (**Table 1**). To increase the number of identified proteins, a typical approach involves even more material and conducts extensive fractionation of the sample. However, numerous biological systems, including biopsies and other rare and precious samples, do not yield such quantities as they only provide hundreds to thousands of cells. Therefore, there remains a need for methods to handle minimal sample inputs with ease, high coverage and reproducibility ^3^.

**Table 1.** The table provides an overview of requirements and expected results for standard proteomics protocols, a recently developed single-cell method, and the minimal input protocol. Rs - Spearman correlation coefficient

|  | **Number of cells prepared** | **Proteins identified** | **Correlation with Standard input sample (Rs)** | **Correlation between replicates (Rs)** | **Correlation between RNA abundance (Rs)** |
| --- | --- | --- | --- | --- | --- |
| **Standard** | 10<sup>6-8</sup> | ~2,700 | - | ~0.99 | ~0.43 |
| **ScoPe-MS** | 1 | ~900 | ~0.62 | ~0.2 | ~0.25 |
| **Minimal input protocol** | 1,000 | ~2,500 | ~0.81 | ~0.99 | ~0.39 |

Recent years have seen several advances in ‘nanoproteomics’, which analyzes small samples from <5,000 cells (corresponding to ∼500 ng proteins) or even single cells^4^. If single cells are used, resulting measurements often suffer from high variation and low reproducibility. The number cells needed depends on the cell type, lysis protocol and measurement method, it is shown to provide enough material for quantitative analysis. For example, using 5,000 human breast cancer cells, previous studies identified 105 to 665 distinct proteins ^5,6^.

Even more recent approaches have tested new technologies and optimized protocols in their use for nanoproteomics. For example, some methods employ specialized systems such as custom designed platforms ^7–9^ that enabled high resolution proteomics in tens of cells or even single cells. Other systems, such as CyTOF ^10^, CITE-seq ^11^ and proximity ligation assays ^12^, have employed specific antibodies, which create sensitive measurements but are limited to proteins with available reagents. Other methods have relied on specific material such as paramagnetic beads ^13^ or collection microreactors ^14^ to maximize yield from little starting material.

A recent nanoproteomics protocol, SCoPE-MS, analyzes proteins from single, hand-picked mammalian cells via Tandem Mass Tagging (TMT) coupled to conventional mass spectrometry ^15^ (**Table 1**). In a standard 10plex TMT experiment, peptides from 10 different samples are labeled with sample-specific mass tags and then pooled. During the subsequent tandem-mass spectrometry experiment, the tags are indistinguishable by mass at the first ‘precursor’ level and therefore isolated together. In the subsequent second level of analysis, the peptides and tags fragment: the peptides can be sequenced and each tag’s channel is quantified through ion intensity measurements (**Table 1**). In SCoPE-MS, one channel in the setup is dedicated to a ‘carrier’ with peptides at high abundance that produce enough signal for reliable peptide identification (**Table 1**). The remaining channels contain the experimental samples. Their peptides’ abundance is too low for identification, but the intensities of the mass tags are available for quantitation. The fundamental idea of the approach is therefore to separate peptide identification and quantitation. While a breakthrough, results from single cells struggle with proteome coverage, reproducibility, and correlation with corresponding transcript abundances (**Table 1**).

To provide a simple protocol with reasonable proteome coverage and reproducibility in cases where more than a single cell is available, we integrated and optimized multiple steps in the proteomics workflow to analyze samples from a thousand cells. The minimal input method uses 1,000 cells, does not require special equipment other than a sonicator for cell lysis and quantifies >2,500 proteins per sample (**Table 1**) with high reproducibility. It can detect two-fold changes in expression levels with statistical significance. The protocol’s reproducibility as well as the correlations with transcript abundances are comparable with those of standard proteomics protocols (**Figure 1a**). We established and validated the method in mouse embryonic stem cells and *in vitro* differentiated motor neurons. We also demonstrated the method’s use in another organism, analyzing the cardiopharyngeal lineage in the tunicate model *Ciona robusta*.

**Figure 1.**
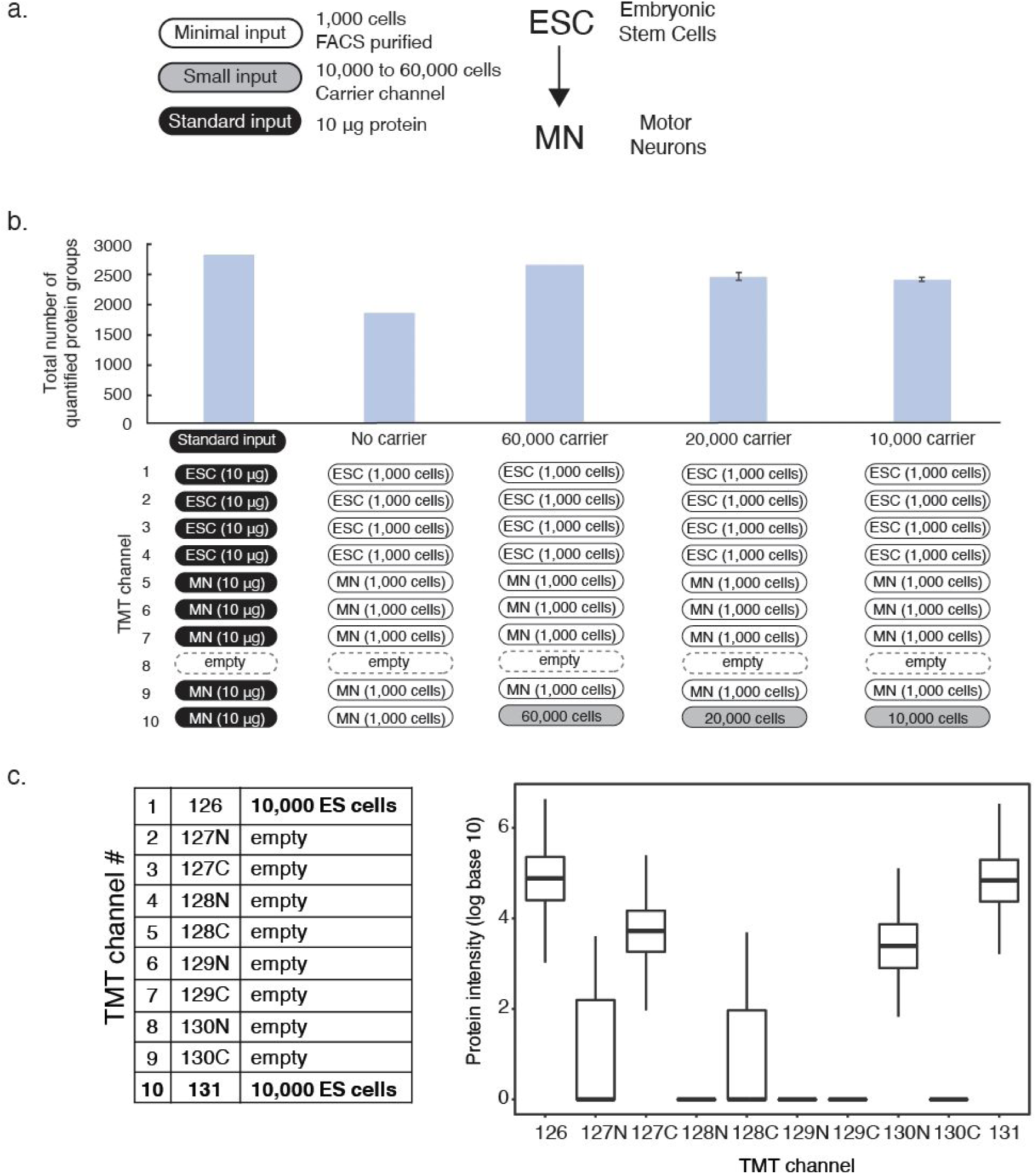
Optimization of the minimal sample protocol. The panels describe testing of the need for a carrier channel, carrier channel input size and position. **a.** The panel shows an overview of the cell numbers used for different purposes in the protocol and comparisons. **b.** We compared TMT setups for using 1,000 cells with and without carrier channel and using 60,000, 20,000, and 10,000 cells for the carrier channel. The graph shows the numbers of protein groups detected using the experimental setup below. The error bars show standard deviation. **c.** We show that the carrier channel affects neighboring channels at 1 thompson distance using a 10plex TMT experiment with empty channels 2-8 and sample from 10,000 cells in channels 1 and 10. ESC - embryonic stem cells, MN - motor neurons, FACS - fluorescence-activated cell sorting, TMT - tandem mass tag

## Experimental Procedures

### Mammalian cell culture

We use the transcription factor cassette NIL (Ngn2-Isl1-Lhx3) to program spinal motor neurons as previously described ^16^. We cultured the mouse embryonic stem cell (ESC) line in 2-inhibitor based medium (Advanced DMEM/F12:Neurobasal (1:1) medium (Thermo Fisher, 12634028, 10888022) supplemented with 2.5% ESC-grade Tet-negative fetal bovine serum (vol/vol, VWR 35–075-CV), 1X N2 (Thermo Fisher, 17502–048), 1X B27 (Thermo Fisher, 17504–044), 2 mM L-glutamine (Thermo Fisher, 25030081), 0.1 mM β-mercaptoethanol (Thermo Fisher, 21985–023), 1000 U/ml leukemia inhibitory factor (Fisher, ESG1107), 3 μM CHIR99021 (Biovision, 1991) and 1 μM PD0325901 (Sigma, PZ0162-5MG), and maintained at 37°C, 8% CO2. We dissociated embryonic stem cells by TrypLE (Gibco, 12605010) and prepared for FACS purification or seeded for differentiation into motor neurons.

To differentiate ESCs into motor neurons, we trypsinized ESCs (Thermo Fisher, 25300–120) and seeded as single cells at 25,000 cells/ml in ANDFK medium (Advanced DMEM/F12: Neurobasal (1:1) Medium (Thermo Fisher, 12634028, 10888022), 10% Knockout SR (vol/vol) (Thermo Fisher, 10828–028), 2 mM l-glutamine (Thermo Fisher, 25030081), and 0.1 mM 2-mercaptoethanol (Thermo Fisher, 21985–023)) to initiate formation of embryoid bodies (EBs) (Day -4) in the suspension culture using 10 cm untreated dishes (Fisher, 08-772-32) and maintained at 37°C, 5% CO2. We changed medium 2 days later (Day -2) with addition of 3 μg/mL Doxycycline (Sigma D9891) to induce the NIL transcription factors. We dissociated the EBs treated with Doxycycline for 2 days, now induced motor neurons, by 0.05% Trypsin-EDTA (Thermo Fisher, 25300–120) and prepared for FACS purification. We estimated the cell counts used for carrier channels using Countess II FL Automated Cell Counter (Thermo Fisher, AMQAF1000).

### *Ciona robusta* handling

Wild *Ciona robusta* were obtained from M-REP (Carlsbad, CA, USA), and kept under constant light to avoid spawning. Gametes from several animals were collected separately for *in vitro* cross-fertilization followed by dechorionation and electroporation as previously described ^17^, and cultured in filtered artificial seawater (FASW) in agarose-coated plastic Petri dishes at 18°C. 50 µg sorting constructs (*Mesp>tagRFP, MyoD905>eGFP* and *Hand-r>tagBFP)* and 70 µg of experimental constructs (*Mesp>LacZ, Mesp>Fgfr*^*DN*^, *Mesp>Mek*^*S216D,S220E*^) were electroporated.

### FACS-purification of mammalian cells

To purify primary motor neurons by fluorescence-activated cell sorting (FACS), we prepared single cell suspensions of each sample to approximately 1–9 × 10^6^ cells/mL. We added 20 uL of 50 ug/mL FDA (Fluorescein Diacetate) to the cell suspension, and isolated motor neurons with a Becton Dickinson ARIA SORP or ARIA II SORP cell sorter. FITC fluorescence was excited with a 488 nm laser and detected with a 530/30 nm (ARIA) or 525/50 nm (ARIA II) filter. We conducted FACS using a ceramic 100 µm nozzle (Becton Dickinson), sheath pressure of 20 pounds per square inch (psi) and a low acquisition rate of 1,000–4,000 events/s. We collected mouse embryonic stem cells and motor neurons in 1.7 ml microcentrifuge tubes containing 20 µl of ice-cold PBS or 0.1% Rapigest (Waters, MA) dissolved in PBS. Total FACS time per experiment was 1–2 h.

### FACS-purification of *Ciona robusta* cells

Sample dissociation and FACS were performed as previously described ^18,19^. Embryos and larvae were harvested at 15 hpf in 5 ml borosilicate glass tubes (Fisher Scientific, Waltham, MA. Cat.No. 14-961-26) and washed with 2 ml calcium- and magnesium-free artificial seawater (CMF-ASW: 449 mM NaCl, 33 mM Na_2_SO_4_, 9 mM KCl, 2.15 mM NaHCO_3_, 10 mM Tris-Cl pH 8.2, 2.5 mM EGTA). Embryos and larvae were dissociated in 2 ml 0.2% trypsin (w/v, Sigma, T-4799) in CMF-ASW by pipetting with glass Pasteur pipettes. The dissociation was stopped by adding 2 ml filtered ice cold 0.05% BSA CMF-ASW. Dissociated cells were passed through 40 μm cell-strainer and collected in 5 ml polystyrene round-bottom tube (Corning Life Sciences, Oneonta, New York. REF 352235). Cells were collected by centrifugation at 800 g for 3 min at 4°C, followed by two washes with ice cold CMF-ASW. Cell suspensions were filtered again through a 40 μm cell-strainer and stored on ice. Cell suspensions were used for sorting within 1 hr. Cardiopharyngeal lineage cells were labeled by *Mesp>tagRFP* reporter. The mesenchyme cells were counter-selected using *MyoD905>GFP* as described ^19^. Dissociated cells were loaded in a BD FACS AriaTM cell sorter. 488 nm laser, FITC filter was used for GFP; 407 nm laser, 561 nm laser, DsRed filter was used for tagRFP and Pacific BlueTM filter was used for tagBFP. The nozzle size was 100 μm.

### Proteome analysis by mass spectrometry

We sonicated cells with Covaris S220 or Diagenode Bioruptor Pico sonicators. Sonication settings employed for 180 s at 125 W power with 10% peak duty cycle for Covaris and 15 cycles 30s on 30s off for Bioruptor, in a degassed water bath at 4 °C. After lysis, we heated the samples for 15 min at 90 °C to denature proteins. We then added 1 ug and 0.5 ug of mass spectrometry grade trypsin (Sigma Aldrich) to standard and minimal samples, respectively, and digested the proteins into peptides at 37°C overnight. We measured resulting peptide concentrations with the Pierce Quantitative Fluorometric Peptide Assay (ThermoFisher) kit for standard samples. We dissolved tandem mass tag (TMT) 10plex reagents (Thermo Scientific) in anhydrous acetonitrile (0.8 mg/82 μl). We labeled peptides per sample with 10 μl of the TMT 10plex label reagent. Following incubation at room temperature for 1 h, we quenched the reactions with 8 μl of 5% hydroxylamine for 15 min. All samples were combined in a new microcentrifuge tubes at equal amounts and reduced to remove acetonitrile using an Eppendorf Concentrator Vacufuge Plus. The salt removal was performed using Pierce™ C18 Spin Tips (Thermo Scientific, #84850) according to the manufacturer’s instructions.

We used an EASY-nLC 1000 coupled on-line to a q-Exactive HF spectrometer (both Thermo Fisher Scientific). Buffer A (0.1% FA in water) and buffer B (80% acetonitrile, 0.5% acetic acid) were used as mobile phases for gradient separation. Separation was performed using a 50 cm x 75 µm i.d. PepMap C18 column (Thermo Fisher Scientific) packed with 2 µm, 100 Å particles and heated at 55°C. We used a 155 min segmented gradient of 0.1% FA (solvent A) and 80% ACN 0.1% FA (solvent B) at a flow rate of 250 nl/min as follows: 2 to 5 %B in 5 min, 5 to 25 %B in 110 min, 25 to 40 %B in 25 min, 49 to 80% B for 5 min and 80 to 95% B for 5min. Solvent B was held at 95% for another 5 min.

For label-free analysis, the full MS scans were acquired with a resolution of 120,000, an AGC target of 3e6, with a maximum ion time of 100 ms, and scan range of 375 to 1500 m/z. Following each full MS scan, data-dependent high-resolution HCD MS/MS spectra were acquired with a resolution of 30,000, AGC target of 2e5, maximum ion time of 150 ms, 1.5 m/z isolation window, fixed first mass of 100 m/z and NCE of 27 with centroid mode. For labeled analysis, the full MS scans were acquired the same settings as label-free. Following each full MS scan, data-dependent high-resolution HCD MS/MS spectra were acquired with a resolution of 60,000, AGC target of 2e5, maximum ion time of 100 ms, 1.2 m/z isolation window, fixed first mass of 100 m/z and NCE of 35 with centroid mode.

### Protein analysis of raw data

The RAW data files were processed using MaxQuant ^20^ (version 1.6.1.0) to identify and quantify protein and peptide abundances. The spectra were matched against the Mus musculus Uniprot database (downloaded August 18, 2018) with standard settings for peptide and protein identification, that allowed for 10 ppm tolerance, a posterior global false discovery rate (FDR) of 1% based on the reverse sequence of the mouse FASTA file, and up to two missed trypsin cleavages. We estimated protein abundance using iBAQ ^21^ for label-free experiments and intensity for TMT experiments. 10plex TMT modifications on Lys and N-terminal amines were considered as fixed modifications. TMT quantification was performed at MS2 level with default mass tolerance and other parameters. We then used the reporter ion intensities as estimates for protein abundance.

### Data analysis

Z-score normalization and Student’s t-test statistics were conducted using Perseus (version 1.5.3.0) ^22^. Student’s t-test analyses were done with permutation based FDR using a cut-off of 0.05 with 250 randomizations. All further computational analyses were conducted in R. Correlation analysis was conducted with the cor() function using raw data. Principal Component Analysis was conducted with the prcomp() function using log base 2 transformed data. RNA-seq data was used from Velasco et al. ^23^ and analyzed using the EdgeR package ^24,25^. Function enrichment analyses were done using Panther 14.1 ^26^. Scatterplots and correlation plots were made using ggplot2 package ^27^.

## Results

### Simplified protein extraction to maximize sample retention

To establish the protocol and assess its performance, we optimized the proteomics workflow at several steps and evaluated proteomic differences in an established *in vitro* differentiation paradigm comparing mouse embryonic stem cells (ESC) and differentiated motor neurons (MN)(**Figure 1a**)^28^. First, we tested different sonicators and buffers for cell lysis using 5,000 mouse embryonic stem cells purified by FACS and label-free mass spectrometry (**Suppl. Figure S1**). We compared the results to those from a standard input sample, which contained protein from ∼500,000 mammalian cells. The standard sample preparation included cell lysis with a phosphate buffered saline-based buffer without detergent, using the Bioruptor sonicator and clean-up of peptides with reverse phase filters. For mass spectrometry analysis, we injected all of the peptides derived from the minimal input samples and 600 ng (∼60,000 cells) from the standard samples.

As we identified similar sumber of peptides and proteins detected in samples from either lysis buffer, we opted for the simpler one, which contained PBS only. In addition, we used the Bioruptor sonicator for the remainder of the experiments as it achieved best protein identification in our hands. However, different samples or sonicator settings might alter this choice.

The results are reproducible with respect to both identified peptides and proteins (**Suppl. Figure 1**). Technical replicates of protein and peptide abundances correlated better than abundances measured across different lysis methods (average R^2^ = 0.93, **Suppl. Figure 2**).We observe higher variation in the number of proteins detected for certain conditions, however this variance does not translate into protein quantification **(Suppl. Figure S2).** For example, even though the number of proteins detected varies in the conditions where Bioruptor was used, the replicates show higher correlation in terms of protein quantification compared to conditions where Covaris sonicator was used. These results were similar in the *Ciona* samples discussed below. We also showed that using samples as small as 1,000 cells in label-free approaches yielded comparable protein identifications and reproducibility to those using 5,000 cells, supporting the use of 5,000 cells for optimization (**Suppl. Figure S3**).

**Figure 2.**
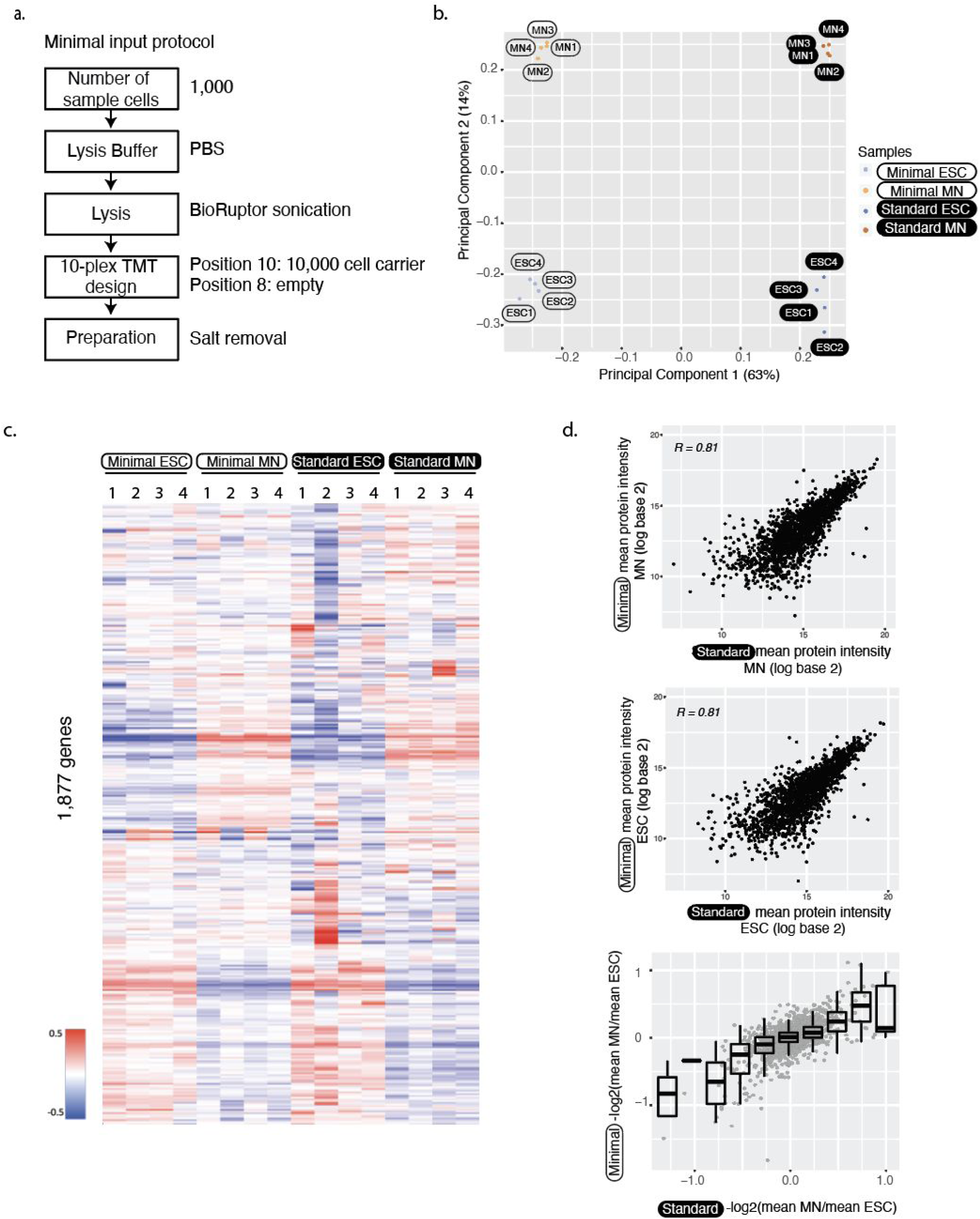
Accurate and reproducible protein abundance measurements. **a.** The flowchart illustrates the minimal input protocol with points of optimization. **b.** The first principal components of minimal and standard sample input data separate both the protocol, but also the two different cell types. The analysis was done using 1,763 protein groups that were identified in both minimal and standard preparations. **c.** Heat map of protein abundances. Abundances were first scaled to the sum of log base 10 equalling 10,000 in each column. Then we removed the first principal component and subtracted the row median from each entry to remove gene-to-gene effect. Note that these normalizations were only used for visualization, not for significance testing for differential expression. We used hierarchical clustering with Manhattan distance measure. Each experiment has 4 replicates. **d.** Protein abundances (top, middle) and fold changes (bottom) from minimal and standard sample input preparations correlate well. The analysis was done using 1,763 protein groups that were identified in both minimal and standard preparations. ESC - embryonic stem cells; MN - motor neurons

### Experimental design for protein quantitation with peptide mass tags

Next, we optimized the use of 10plex TMT labeling with a carrier channel. We used a pool of equal proportions of both cell types analyzed in the experimental samples as a carrier. First, we showed that a setup with a carrier channel outperformed a setup without a carrier, i.e. with minimal input in all channels, with respect to the number of detected proteins (“No carrier”, **Figure 1c**, first 3 bars). Without a carrier channel, the amount (ng) of protein contributing to the peptide signal is derived from 9 x 1,000 = 9,000 cells, while for the setup with a carrier channel, the sample is derived from at least 10,000 + 8 x 1,000 = 18,000 cells (or more if the carrier is larger). The results suggest that samples from 9,000 cells do not provide enough material for maximal peptide identification, at least not from comparatively small cells used here such as embryonic stem cells and motor neurons.

Second, we showed that the carrier should ideally be placed in channel 10, leaving channel 8 empty. The reason for this setup lies in the carrier channel producing signals ‘spilling over’ into channel 8 as we show in **Figure 1d**. In **Figure 1d**’s setup, all TMT channels are empty except for positions 1 and 10 which both contained peptides prepared from 10,000 cells. The carrier in channel 1 produced signals in channels 2, 3 and 5. The carrier in channel 10 did not affect channel 9 immediately adjacent to it. However, we observed a substantial signal at the -1 Thompson distance to the carrier in channel 10, i.e. in channel 8, indicating contamination of the mass tag with the light isotope. Indeed, the observed intensity in channel 8 was about 3-4% of the total intensity in channel 10, consistent with the contamination with the light isotope as reported by the manufacturer. For this reason, we placed the carrier into channel 10 and left channel 8 empty in all subsequent TMT experiments. This setup should be robust even if the reported impurity of the channel 10 label differs.

Finally, we minimized the size of the carrier channel, as a low carrier-to-sample ratio is advantageous with respect to highest reproducibility (**Suppl. Figure S4**). We tested carrier channels with peptides derived from 60,000, 20,000, and 10,000 cells. We found that the carrier from 10,000 cells provided similar protein identification and reproducibility to the larger carrier channels. We thus used this carrier size in subsequent experiments, resulting in a final carrier-to-individual sample ratio of 10,000:1,000 = 10:1 (**Figure 1b**, last 3 bars).

### Validating accuracy of measured protein abundance

**Figure 2a** shows an overview of the final protocol which uses 1,000 cells in experimental sample channels and 10,000 cells in the carrier channel. We then evaluated this workflow with respect to coverage and reproducibility of the measured protein abundance. To do so, we used an established *in vitro* motor neuron differentiation protocol for mouse embryonic stem cells ^28^. While requiring only 1/50 of the number of cells, the minimal input protocol’s proteome coverage reached 88% of what we detected in single-shot standard proteome analysis, i.e. 2,483 compared to 2,828 proteins (**Figure 1c**). Replicate experiments in the minimal input setup correlated with an average of R^2^=0.99 in log-log abundance plots (**Suppl. Figure S5-S7**), indicating high reproducibility.

Next, we used principal component analysis to compare results from the standard and minimal input preparations (**Figure 2b**). As expected, the first component separated the two experimental setups which differed as described above, e.g. through use or omission of FACS-purification. However, the second component separated the two cell types, indicating that the protocol is able to produce biologically meaningful protein quantitation. Indeed, when removing the first principal component, we observe striking similarities between samples from the minimal and standard input preparations (**Figure 2c**). Both preparations show similar differences between undifferentiated and differentiated cells.

We further confirmed the consistency between the minimal input and the standard protocol by direct correlation of the measured protein abundances in the two cell types: the Pearson correlation coefficient ranged between 0.81 and 0.77 between the two protocols (**Figure 2d, Suppl. Figure S6-S7**). Both minimal input and standard input preparations also showed similar correlation with corresponding transcript abundances as taken from bulk RNA sequencing from the same differentiation paradigm ^23^, with Spearman correlation coefficients ranging from 0.39 to 0.43 (**Suppl. Figure S8**).

To further validate the protocol’s ability to identify differentially expressed proteins, we turned to known markers of successful differentiation, which we used as positive controls. Several key neuron marker genes, e.g. AINX ^29^, MAP1B ^30^, RABP1 ^30,31^, STMN2 ^32^ and TBB3 ^33^, were up-regulated in MN cells (**Figure 3a**). The differentially expressed proteins were also significantly enriched in plasma membrane proteins (q-value<0.01), including the known neuron markers MAP1B and STMN2.

**Figure 3.**
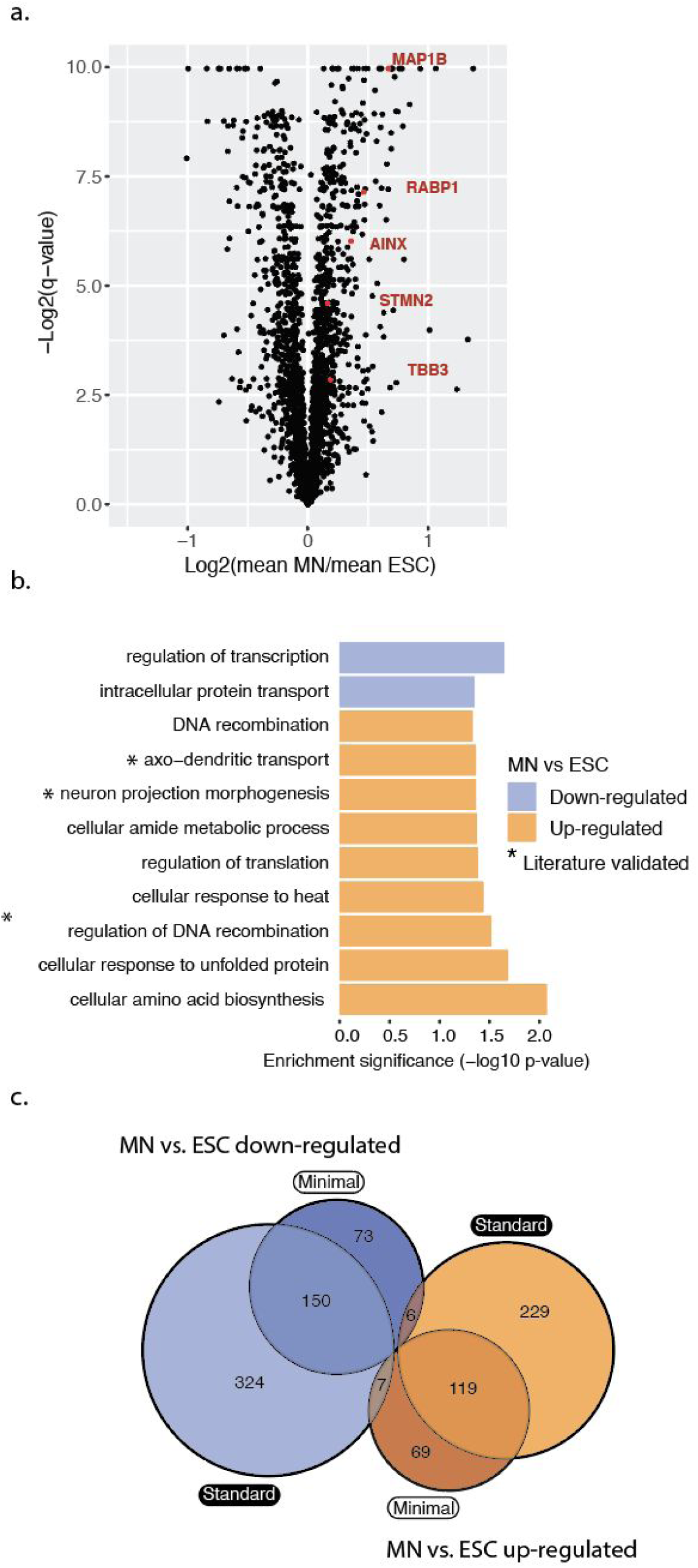
Proteins differentially expressed between ESC and MN. a. Volcano plot showing in red widely recognized neuronal markers over-expressed in MNs compared to ESCs in the minimal input preparation. q-values are calculated using Student’s t-test, using permutation based FDR with a cut-off of 0.05. **b.** Differentially expressed proteins have similar function enrichments in both the minimal and standard input protocol (p-value<0.05). **c.** The minimal and standard input protocols overlap in their results with respect to identification of differentially expressed proteins: significantly up- or down-regulated proteins (MN vs. ESC) from either preparation overlap with p = 2.070e^-192^ and p = 7.639e^-164^, respectively (hypergeometric test). There are close to no proteins shared across opposing groups (p = 0.221 and p = 0.191, hypergeometric test). ESC - embryonic stem cells; MN - motor neurons

We then tested variation of protein abundances between undifferentiated and differentiated cells against variation between replicates, using the Student’s t-test. The approach identified 229 and 195 proteins with significant up- and down-regulation in motor neurons compared to stem cells, respectively (q-value<0.01, **Figure 3a**). The proteins up-regulated in motor neurons were significantly enriched in several functions such as axo-dendritic transport and neuron projection morphogenesis (p-value<0.05, Fisher’s exact test, **Figure 3b, Suppl. Figure S9**). We compared the significantly differentially expressed proteins between the minimal input and standard sample preparations (**Figure 3c**). While standard preparation identified more differentially expressed proteins due to lower variability across replicates (**Suppl. Figure 10**), the up- and down-regulated proteins from either preparation overlapped significantly (p<0.0001 for MN vs. ESC down-regulated and up-regulated, hypergeometric test). There were virtually no proteins in the opposing groups (p=0.22 and p=0.19, hypergeometric test).

Due to the semi-stochastic nature of proteomics experiments, the minimal input protocol identified some differentially expressed proteins that were missed by the standard input preparation. The proteins specific to the minimal input protocol had abundances ranging over 7 orders of magnitude (**Suppl. Figure S11**), illustrating that abundance alone does not account for preparation specific identification. The minimal input-specific proteins included EZRI, PEBP1 and TBB5 which were up-regulated in motor neurons and which all have known roles in neuronal cells. Combined, these results support the protocol’s ability to quantify proteins and to identify significantly differentially expressed proteins ^34^.

### Applying the minimal input protocol to sorted *Ciona robusta* cells

To test whether the protocol is generalizable to different systems, we used the model chordate *Ciona robusta*. We FACS-purified the cardiopharyngeal lineage using an established protocol ^19^ and applied our method using two different setups. The first experiment employed 1,000 cells for each experimental channel with 4 carrier channels **(Suppl. Figure S12a).** It identified 1,904 proteins. The carrier channels consisted of whole embryo cells, since collecting more than a few thousand cells per condition from the cardiopharyngeal lineage is not feasible. In the second experiment, we omitted the carrier channel and used 5,000 cells for each experimental condition, which resulted in 732 identified proteins.

Next, we assessed reproducibility between biological replicates (**Suppl. Figure S12b**,**c**). Both experiments show reasonable correlation between replicates of log transformed protein abundances, slightly lower than the correlation between replicates of log transformed transcript abundances (**Suppl. Figure S13a**,**b**). When comparing RNA and protein abundances directly, we observed small, but statistically significant positive correlations (p-value < 0.05; **Suppl. Figure S14a**,**b**), indicating that indeed, the minimal input protocol provides meaningful protein abundance estimates.

In sum, the reproducibility and coverage of the *Ciona* experiments are comparable with the results that we have obtained with the mammalian cells **(Suppl. Figure S5, Suppl. Figure S12bc)**. However, we detect no statistically significant differentially expressed genes between conditions. While variation between replicates might contribute to this result, we also speculate that the experimental setup did not allow for detectable proteome changes: the experiment was performed 15 hours post fertilization, about an hour after the cells were born, which is enough time to change the transcriptome ^35,36^, but not enough to translate large numbers of new proteins.

## Discussion

We present a straightforward protocol for proteomics analysis of minimal input samples, i.e. 1,000 cells. The method quantifies ∼2,500 proteins in mammalian samples and sensitively identifies differential expression. Protein abundances measured by the minimal input protocol correlate well with corresponding transcript abundances, and have high reproducibility across replicates. This result is consistent with what is known about the relationship between protein and RNA^34^, and confirms the validity of our measurements. Therefore, the minimal input protocol offers an alternative to existing small input methods, such as SP3 ^13^ or in-StageTip digestion ^37^. SP3 can handle samples from as few as 1,000 cells, however the authors indicate that this amount approached the limit of sensitivity and skewed the quantification^13^. Further, SP3 method requires specialized equipment, such as paramagnetic beads. The in-StageTip method can handle 1 µg of protein ^38^, which corresponds to roughly 10,000 cells, i.e. 10-times more than the protocol presented here.

Our protocol uses sample amounts achievable by, for example, dissection of specific cell types or tissues *in vivo* or FACS purification of rare cell populations, enabling analysis of highly specific cell populations. The minimal input protocol relies on use of a carrier channel, as pioneered for single-cell analysis ^15^. We also tested the protocol for cases in which obtaining a carrier channel is not feasible. We find that the protocol works, in principle, but accuracy of identification and protein quantitation is lower. With a carrier channel, the minimal input protocol achieves reproducibility similar to that of standard proteomics. In addition, the protocol does not require specific equipment or reagents other than anything you can find in a mass spectrometry lab. Therefore, it enables analysis of systems in which it is very difficult to obtain large numbers of cells.

## Acknowledgements

B.V. acknowledges funding by American Heart Association grant #18PRE33990254. D.E.I. acknowledges funding by the NIH/NINDS (1F31NS103447-01A1). B.V. and D.E.I. acknowledge Fleur Strand Graduate Fellowship. E.O.M acknowledges funding by Project ALS (A13-0416). L.C. acknowledges funding by NIH/NHLBI R01 HL108643 and by the Leducq Foundation Trans-Atlantic network of excellence award 15CVD01. C.V. acknowledges funding by the NIH/NIGMS (R35GM127089), and the Zegar Family Foundation Fund for Genomics Research at New York University. We thank Judit Villen and Matthias Selbach for productive discussions.

## Data availability

The mass spectrometry data including the MaxQuant output files have been deposited to the ProteomeXchange Consortium via the PRIDE ^39^ partner repository with the dataset identifier PXD015123.

