## Supplemental for "A simple method to quantify protein abundances from one thousand cells"

**Figure S1:** We optimized the buffer composition and cell lysis method; the graph shows the numbers of **a**. Protein groups and **b**. peptides detected by label-free mass spectrometry using 5,000 embryonic stem cells. The error bars show standard variation.

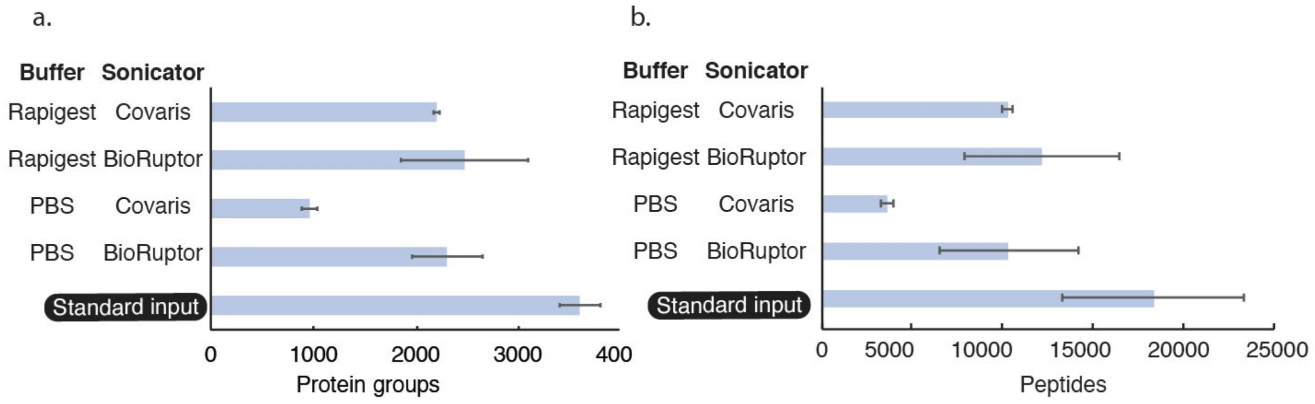


**Figure S2.** Different sample preparation techniques correlate between each other and are highly reproducible (Spearman correlation coefficient). ESC - embryonic stem cells, MN - motor neurons

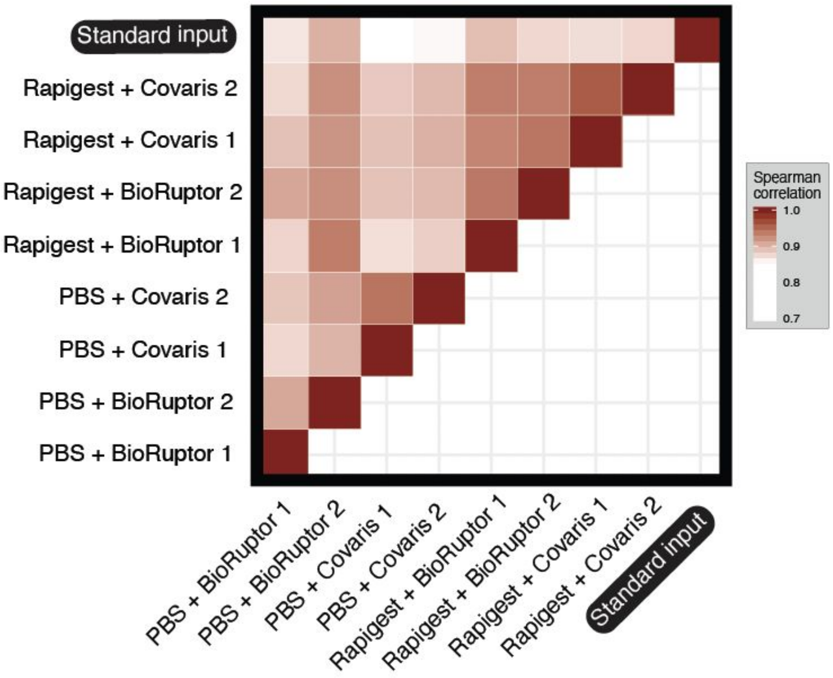

**Figure S3.** We compared the number of protein groups identified using 1,000 and 5,000 embryonic stem cells using label-free quantification and observed no significant differences.

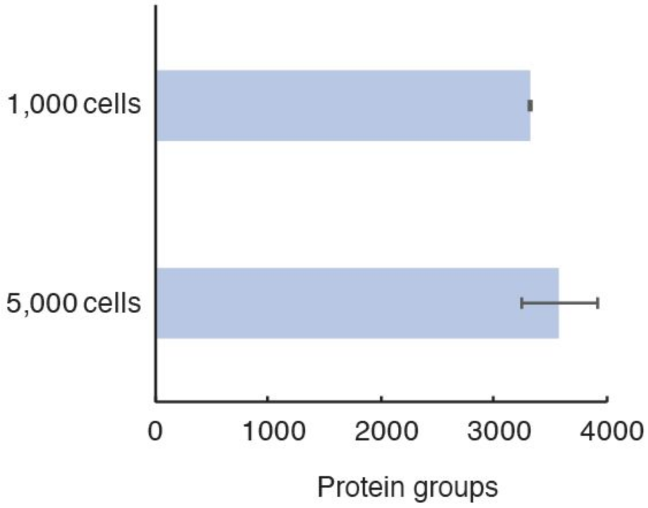


**Figure S4.** We optimized the size of the carrier channel, and to do so we used samples from 60,000, 20,000, and 10,000 cells as carrier channels with 1,000 experimental channels. The graphs show that 10,000 carrier channel shows the most consistent quantification (Spearman correlation) at similar numbers of total protein identification (see main text). ESC - embryonic stem cells, MN - motor neurons

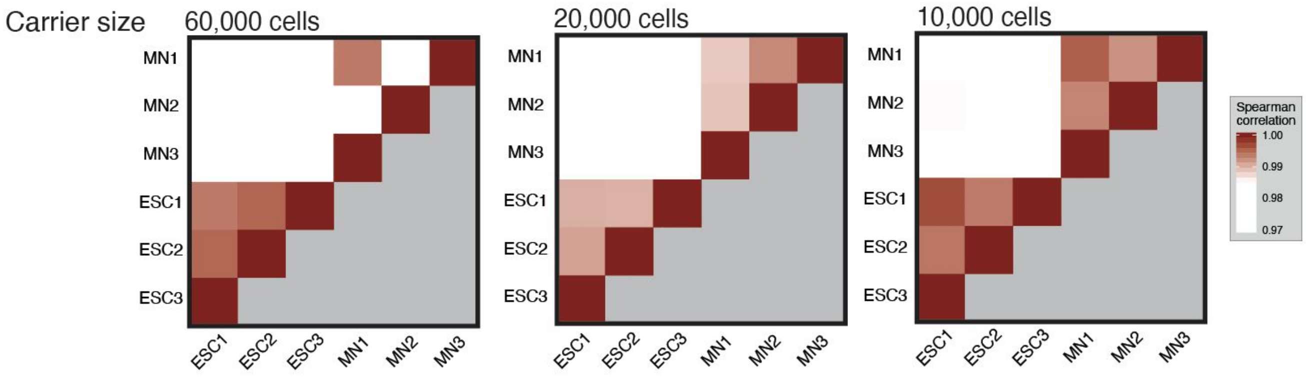

**Figure S5.** The replicates of minimal input MN (left) and ESC (right) show very high correlations of Spearman $R^2$=0.99 and $R^2$=0.98, respectively. ESC - embryonic stem cells, MN - motor neurons

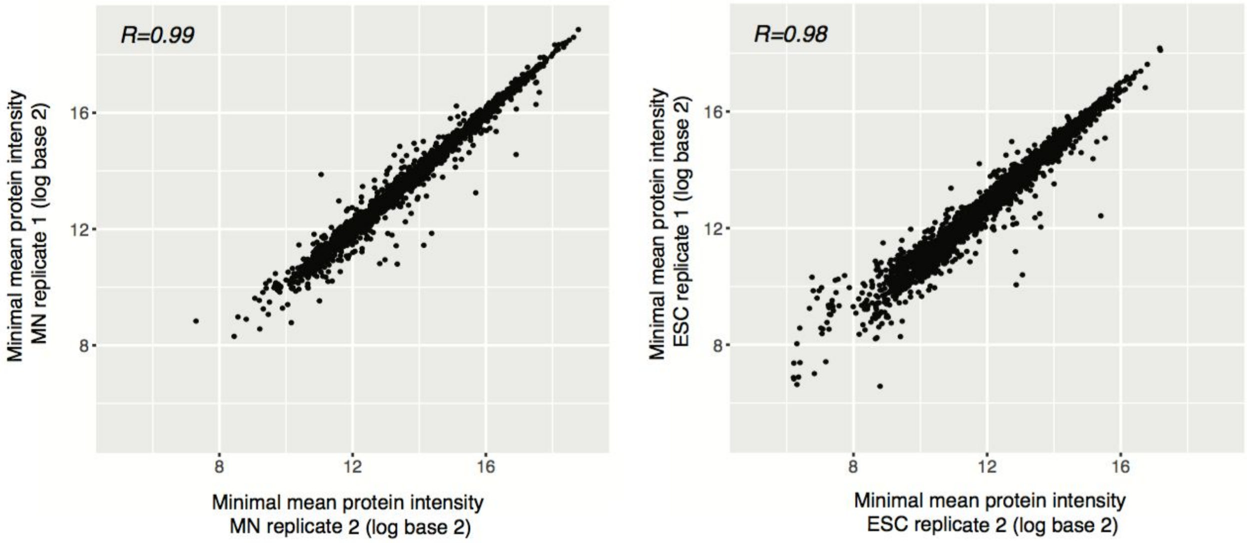

**Figure S6.** Correlations coefficients (Spearman) of quadruplicates resulting from minimal and standard input embryonic stem cells (ESCs).

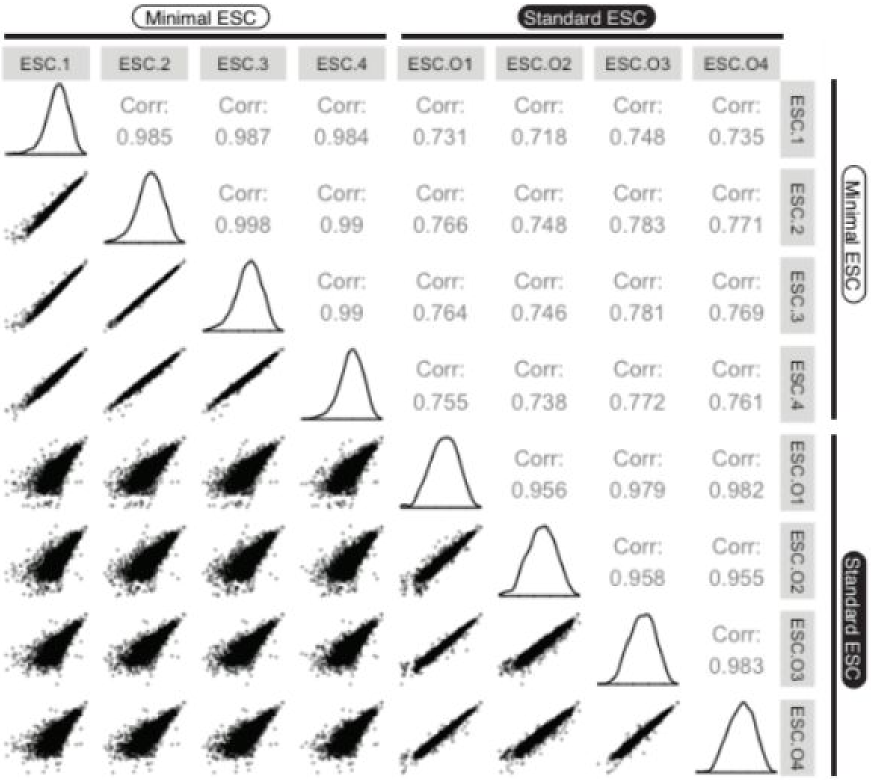

**Figure S7.** Correlations coefficients (Spearman) of quadruplicates resulting from minimal and standard input motor neurons (MNs).

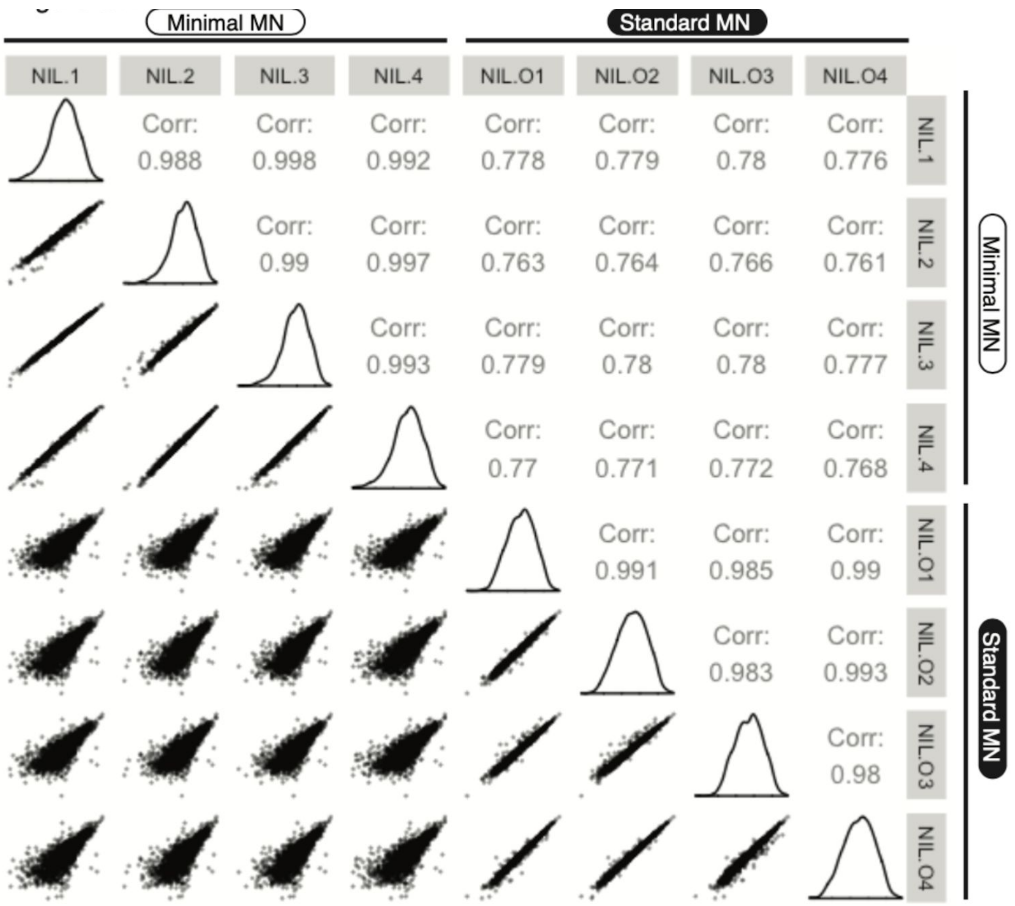

**Figure S8.** Minimal (top) and standard (bottom) input preparations show similar correlation with corresponding transcript abundances, with Spearman coefficients ranging from 0.39 to 0.43. ESC - embryonic stem cells, MN - motor neurons

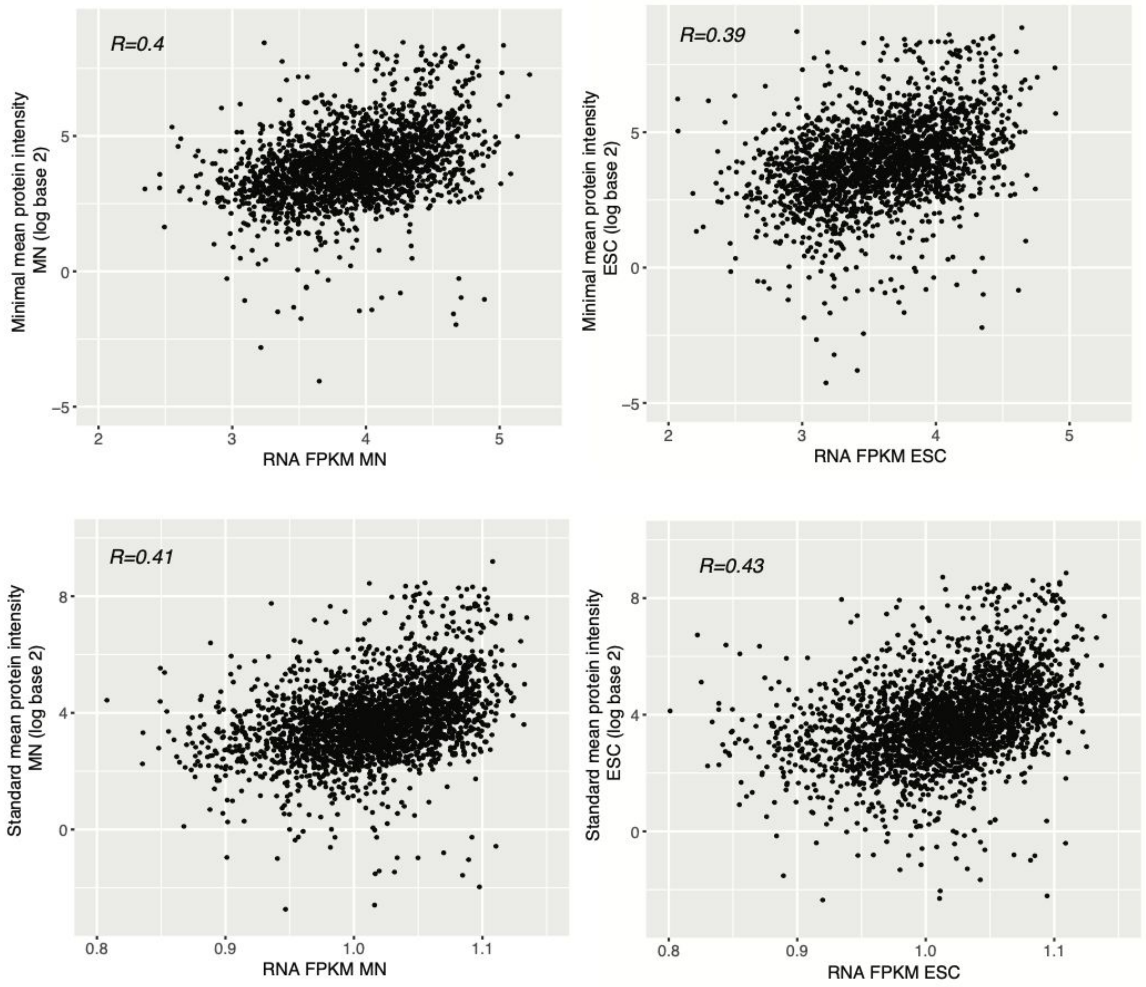

**Figure S9.** Differentially expressed proteins have similar function enrichments between the minimal input and standard protocol (p-value<0.05). ESC - embryonic stem cells, MN - motor neurons

Figure S9.

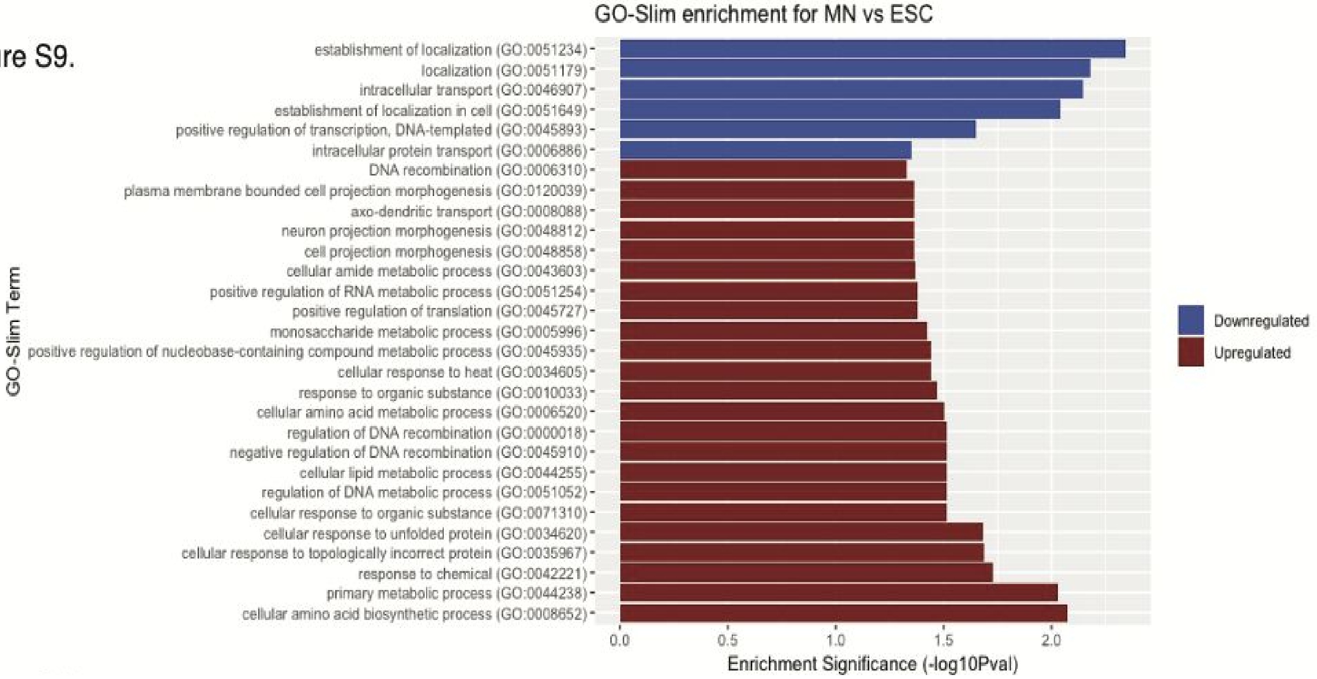

**Figure S10.** Volcanos plot showing in red the significantly different genes in experimental conditions q-value<0.05. Student t-test was used, permutation based FDR with cut-off 0.05. ESC - embryonic stem cells, MN - motor neurons

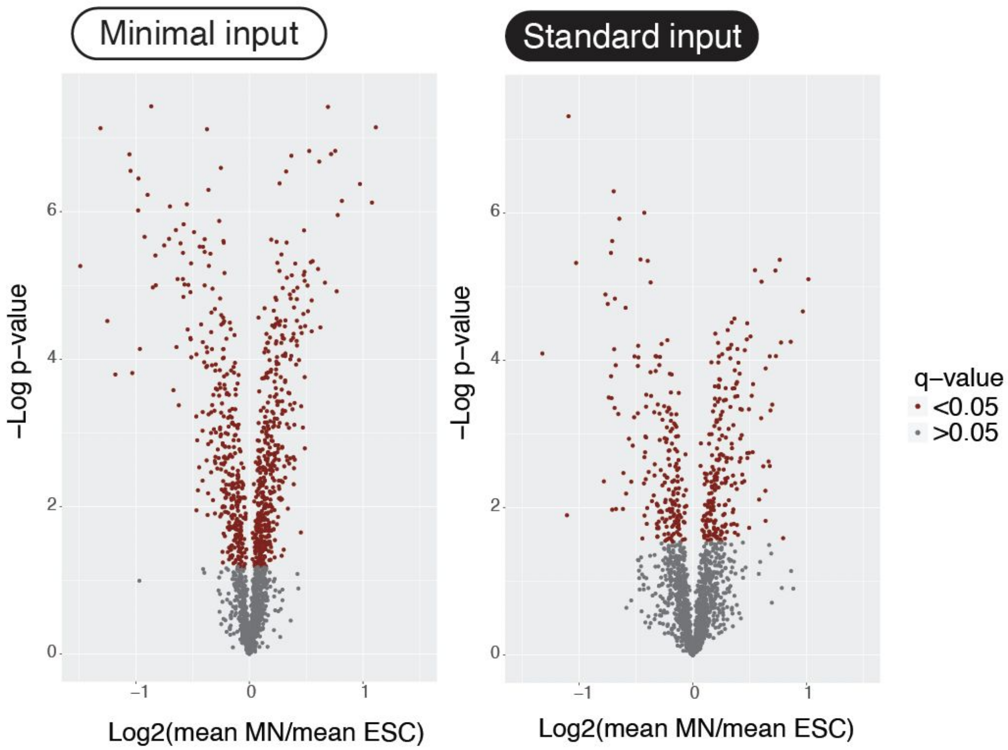

**Figure S11.** Differentially expressed proteins identified by the minimal input preparation agree with results from standard input preparations. The differentially expressed protein groups from the minimal input data is plotted as a volcano plot (top) and an intensity scatter plot (bottom) showing the protein groups that are specific to the minimal input prep in grey, identified and significantly differential in both minimal and standard input preparations in red and not significantly differentially expressed in the minimal input preparation in blue. Significant in both: 552, Significant in minimal and not detected in standard: 265, Significant in minimal and not significant in standard: 135. ESC - embryonic stem cells, MN - motor neurons

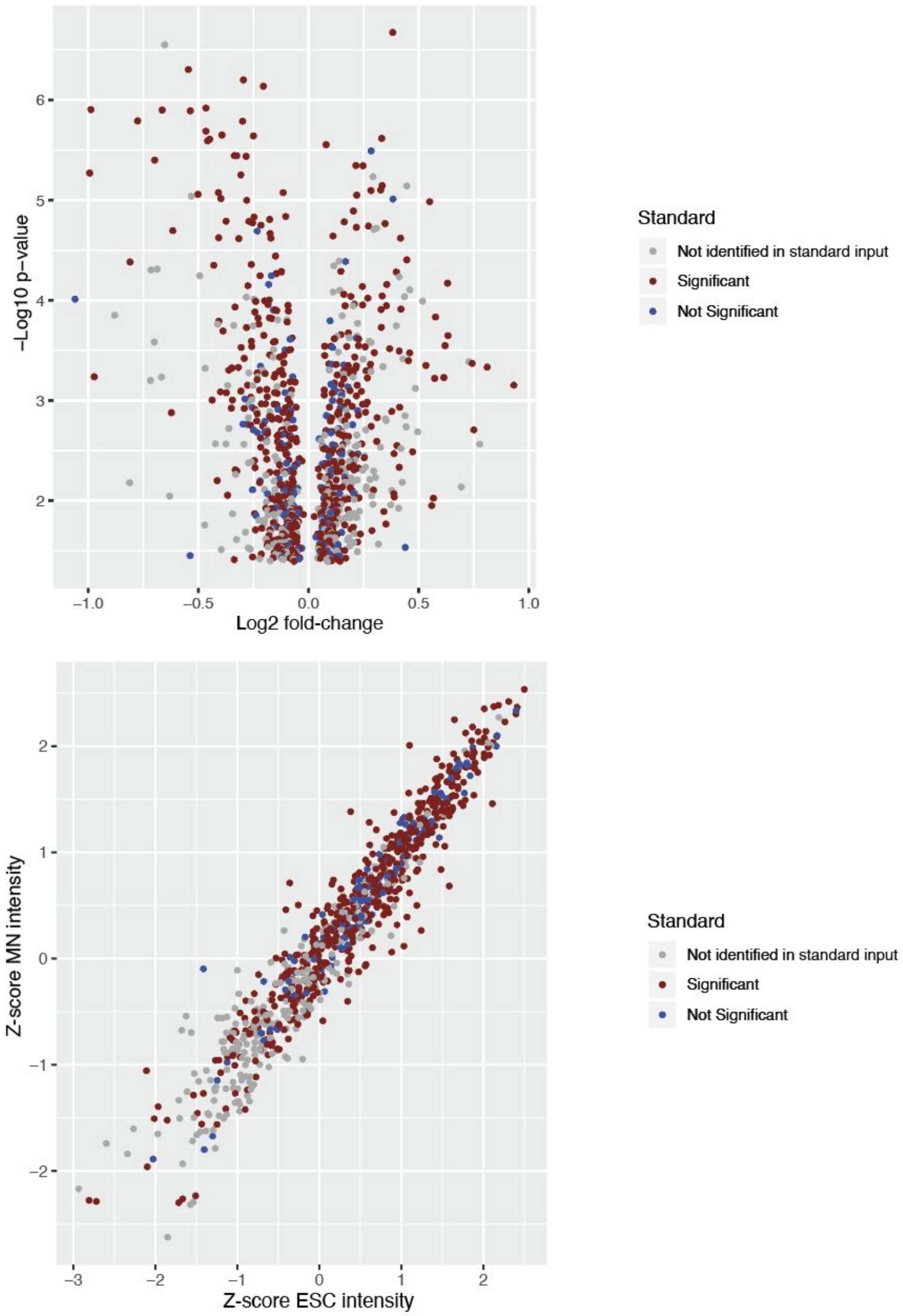

**Figure S12.** Minimal input protocol using sorted *Ciona robusta* cardiopharyngeal lineage. **a.** We tested two experimental designs. Experiment 1 used 1,000 cells for each experimental channel with 4 carrier channels of 20,000 whole embryo cells. It identified 1,904 proteins. Experiment 2 used 5,000 cells for each experimental condition with leaving channel 8 empty. It identified 732 identified proteins. **b.** Experiment 1: scatter plots show reproducibility between protein intensity measurements across replicates. Values indicate Pearson correlation coefficients, ranging from 0.949 to 0.978. **c.** Experiment 2: scatter plots show reproducibility between protein intensity measurements across replicates. Values indicate Pearson correlation coefficients, ranging from 0.946 to 0.984. ASM: Atrial Siphon Muscle, condition electroporated *Mesp>Mek*$^{S216D,S220E}$. Heart: condition electroporated *Mesp>Fgfr*$^{DN}$. Mixed: condition electroporated *Mesp>LacZ*.

**Figure S12a.**

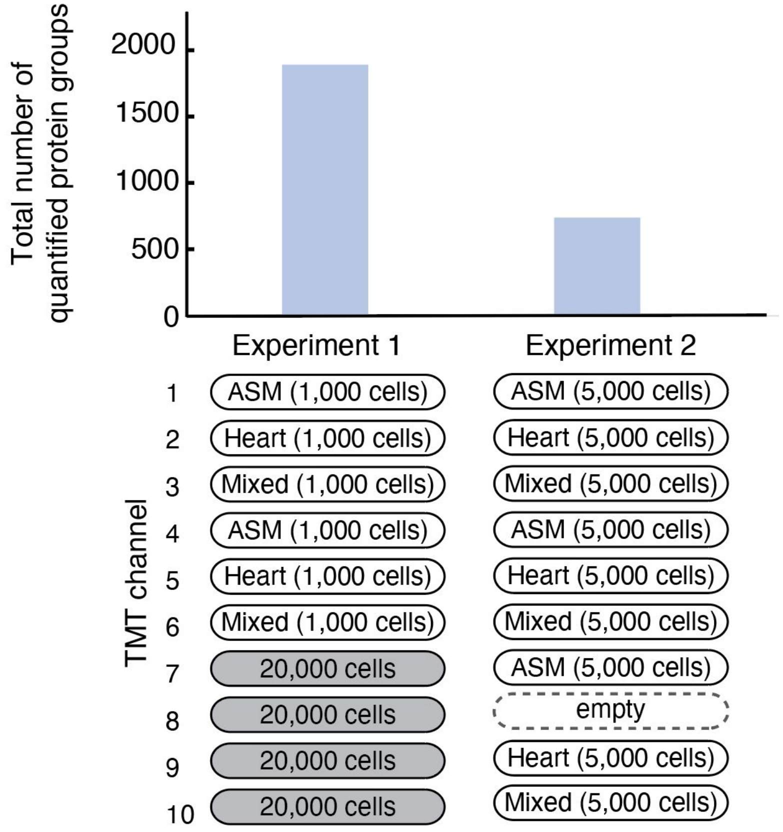

**Figure S12b.**

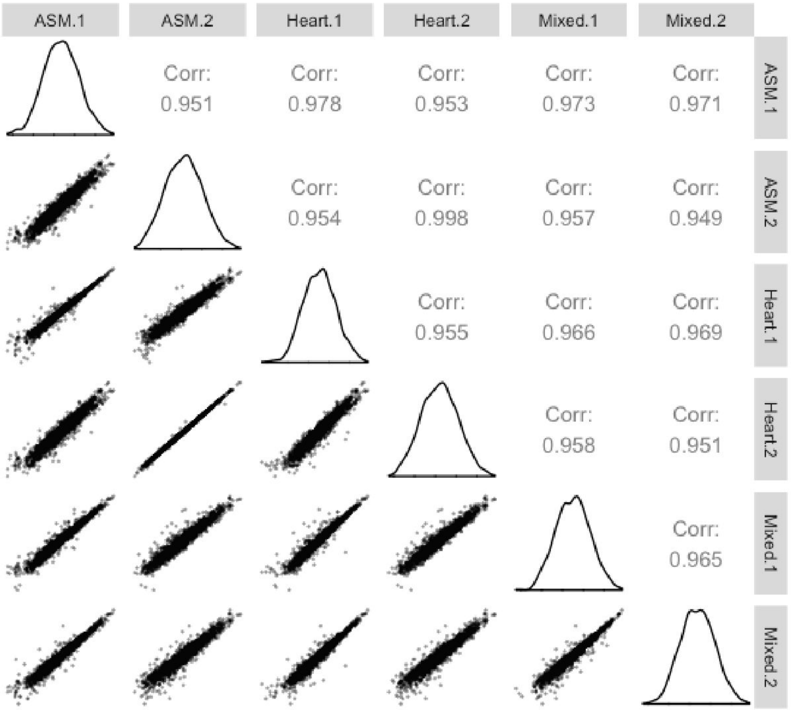


**Figure S12c.**

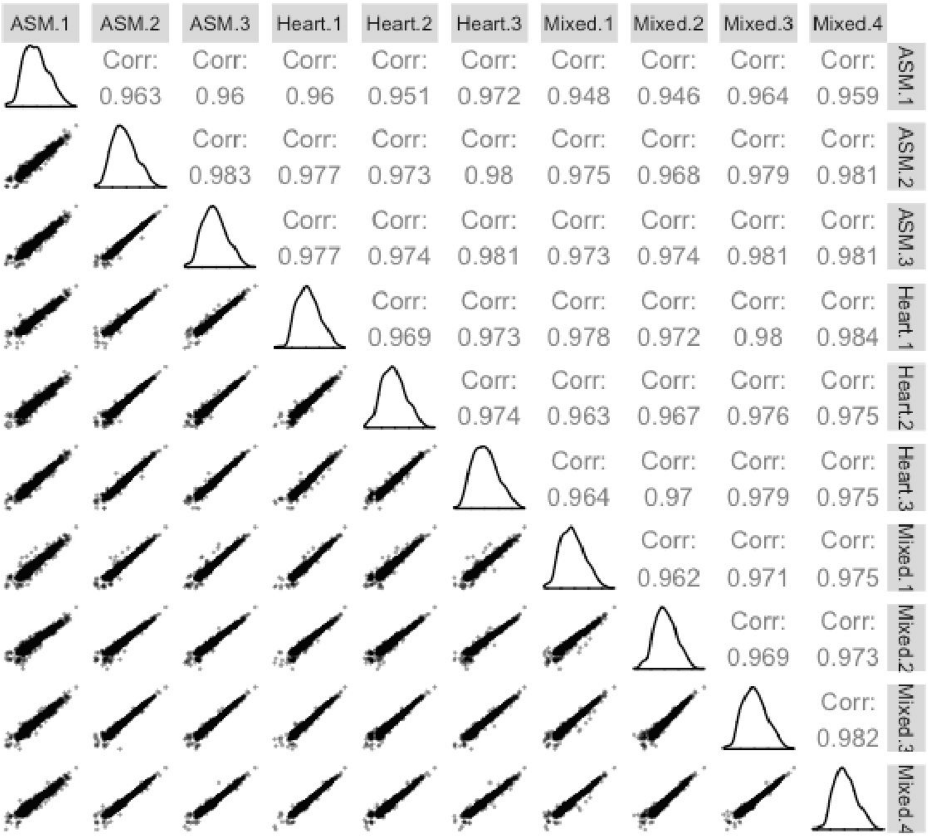

**Figure S13. a.** Experiment 1: scatter plots show reproducibility between RNA count data measurements across replicates. Values indicate Pearson correlation coefficients, ranging from 0.931 to 0.986. **b.** Experiment 2: scatter plots show reproducibility between RNA count data measurements across replicates. Values indicate Pearson correlation coefficients, ranging from 0.944 to 0.99. ASM: Atrial Siphon Muscle, condition electroporated *Mesp>Mek*$^{S216D,S220E}$. Heart: condition electroporated *Mesp>Fgfr*$^{DN}$. Mixed: condition electroporated *Mesp>LacZ*.

**Figure S13a.**

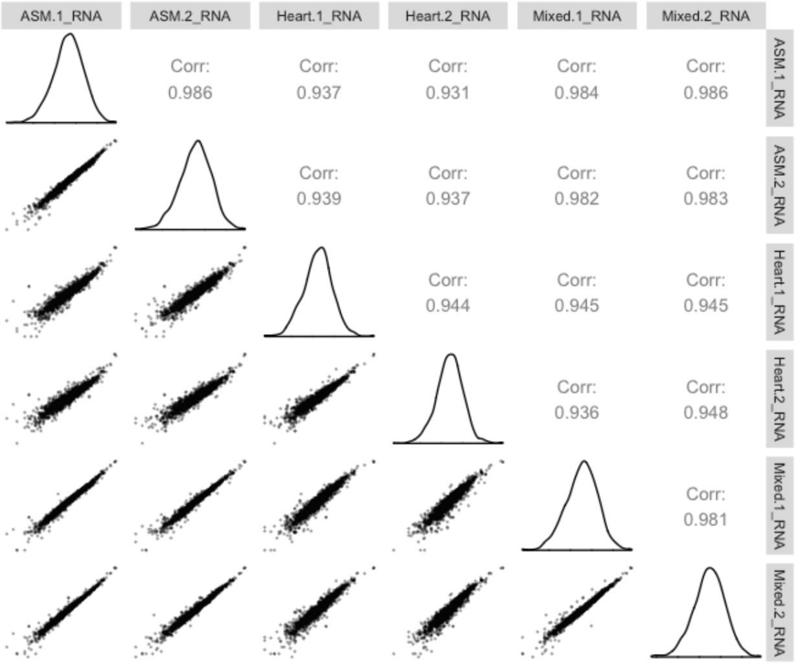


**Figure S13b.**

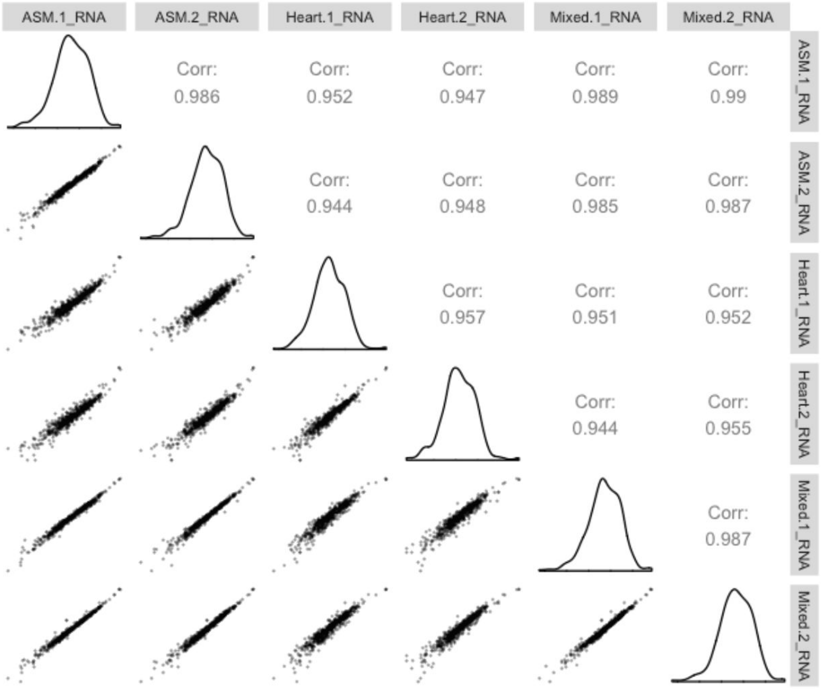

**Figure S14. a.** Experiment 1: The scatter plots show the correlation between log base 2 transformed RNA and protein abundances in measurements from *Ciona robusta* cells. Values indicate Spearman correlation coefficients. All correlations are significant at a p-value<0.01. **b.** Experiment 2: The scatter plots show the correlation between log base 2 transformed RNA (x-axis) and protein abundances (y-axis) in measurements in *Ciona* cells. Values indicate Spearman correlation coefficients. ASM: Atrial Siphon Muscle, condition electroporated *Mesp>Mek*$^{S216D,S220E}$. Heart: condition electroporated *Mesp>Fgfr*$^{DN}$. Mixed: condition electroporated *Mesp>LacZ*.

**Figure S14a.**

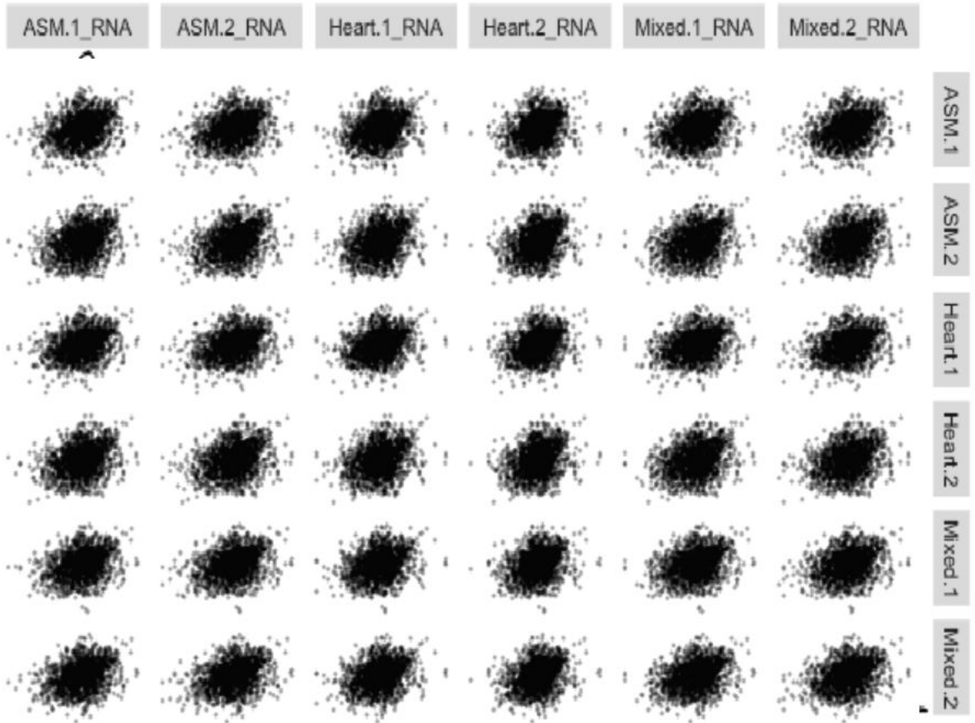


| | ASM.1_RNA | ASM.2_RNA | Heart.1_RNA | Heart.2_RNA | Mixed.1_RNA | Mixed.2_RNA |
|---|---|---|---|---|---|---|
| ASM.1 | Corr: 0.277 | Corr: 0.295 | Corr: 0.276 | Corr: 0.295 | Corr: 0.288 | Corr: 0.292 |
| ASM.2 | Corr: 0.279 | Corr: 0.295 | Corr: 0.277 | Corr: 0.296 | Corr: 0.29 | Corr: 0.294 |
| Heart.1 | Corr: 0.258 | Corr: 0.28 | Corr: 0.256 | Corr: 0.28 | Corr: 0.271 | Corr: 0.273 |
| Heart.2 | Corr: 0.248 | Corr: 0.277 | Corr: 0.248 | Corr: 0.277 | Corr: 0.267 | Corr: 0.265 |
| Mixed.1 | Corr: 0.27 | Corr: 0.286 | Corr: 0.268 | Corr: 0.287 | Corr: 0.283 | Corr: 0.283 |
| Mixed.2 | Corr: 0.272 | Corr: 0.294 | Corr: 0.271 | Corr: 0.295 | Corr: 0.285 | Corr: 0.288 |

**Figure S14b.**

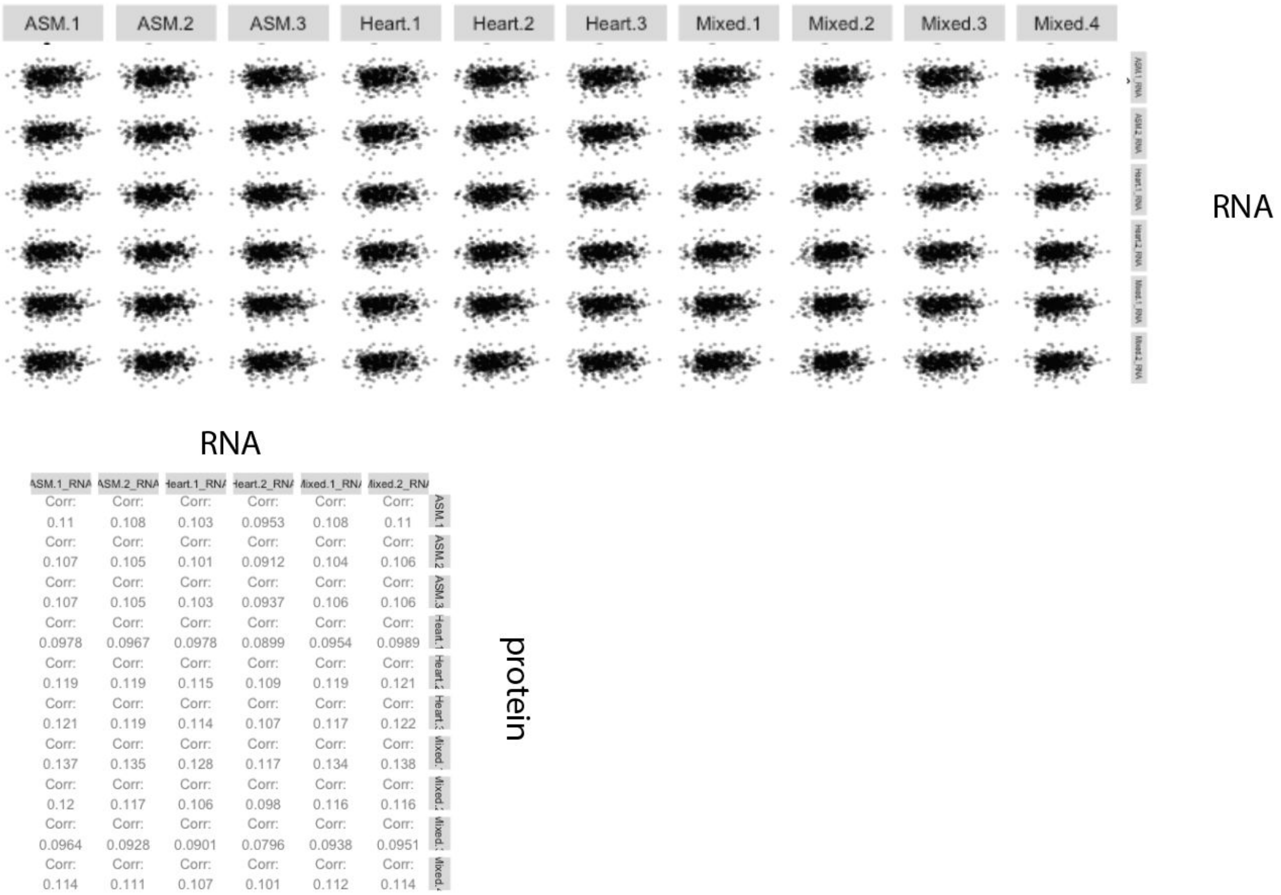
ASM.1 ASM.2 ASM.3 Heart.1 Heart.2 Heart.3 Mixed.1 Mixed.2 Mixed.3 Mixed.4
RNA
RNA
ASM.1_RNA ASM.2_RNA Heart.1_RNA Heart.2_RNA Mixed.1_RNA Mixed.2_RNA
ASM.1: Corr: 0.11 Corr: 0.108 Corr: 0.103 Corr: 0.0953 Corr: 0.108 Corr: 0.11
ASM.2: Corr: 0.107 Corr: 0.105 Corr: 0.101 Corr: 0.0912 Corr: 0.104 Corr: 0.106
ASM.3: Corr: 0.107 Corr: 0.105 Corr: 0.103 Corr: 0.0937 Corr: 0.106 Corr: 0.106
Heart.1: Corr: 0.0978 Corr: 0.0967 Corr: 0.0978 Corr: 0.0899 Corr: 0.0954 Corr: 0.0989
Heart.2: Corr: 0.119 Corr: 0.119 Corr: 0.115 Corr: 0.109 Corr: 0.119 Corr: 0.121
Heart.3: Corr: 0.121 Corr: 0.119 Corr: 0.114 Corr: 0.107 Corr: 0.117 Corr: 0.122
Mixed.1: Corr: 0.137 Corr: 0.135 Corr: 0.128 Corr: 0.117 Corr: 0.134 Corr: 0.138
Mixed.2: Corr: 0.12 Corr: 0.117 Corr: 0.106 Corr: 0.098 Corr: 0.116 Corr: 0.116
Mixed.3: Corr: 0.0964 Corr: 0.0928 Corr: 0.0901 Corr: 0.0796 Corr: 0.0938 Corr: 0.0951
Mixed.4: Corr: 0.114 Corr: 0.111 Corr: 0.107 Corr: 0.101 Corr: 0.112 Corr: 0.114
protein
